# Rhinovirus reduces the severity of subsequent respiratory viral infections by interferon-dependent and -independent mechanisms

**DOI:** 10.1101/2020.11.06.371005

**Authors:** James T. Van Leuven, Andres J. Gonzalez, Emmanuel C. Ijezie, Alexander Q. Wixom, John L. Clary, Maricris N. Naranjo, Benjamin J. Ridenhour, Craig R. Miller, Tanya A. Miura

## Abstract

Coinfection by unrelated viruses in the respiratory tract is common and can result in changes in disease severity compared to infection by individual virus strains. We have previously shown that inoculation of mice with rhinovirus (RV) two days prior to inoculation with a lethal dose of influenza A virus (PR8), provides complete protection against mortality and reduces morbidity. In this study, we extended that finding to a second lethal respiratory virus, pneumonia virus of mice (PVM) and analyzed potential mechanisms whereby RV reduces lethal viral pneumonia caused by PR8 and PVM. RV prevented mortality and weight loss associated with PVM infection, suggesting that RV-mediated protection is more effective against PVM than PR8. Major changes in host gene expression upon PVM infection were delayed compared to PR8, which likely provides a larger time frame for RV-induced gene expression to alter the course of disease. Overall, RV induced earlier recruitment of inflammatory cells, while these populations were reduced at later times in RV-inoculated mice. Findings common to both virus pairs included upregulated expression of mucin-associated genes and dampening of inflammation-related genes in mice that were inoculated with RV prior to lethal virus infection. However, type I IFN signaling was required for RV-mediated protection against PR8, but not PVM. IFN signaling had minor effects on PR8 replication and contributed to controlling neutrophilic inflammation and subsequent hemorrhagic lung pathology in RV/PR8 infected mice. These findings, combined with differences in virus replication levels and disease severity, suggest that the suppression of inflammation in RV/PVM infected mice may be due to early, IFN-independent suppression of viral replication, while in RV/PR8 infected mice may be due to IFN-dependent modulation of immune responses. Thus, a mild upper respiratory viral infection can reduce the severity of a subsequent severe viral infection in the lungs through virus-dependent mechanisms.

**Author Summary:** Respiratory viruses from diverse families co-circulate in human populations and are frequently detected within the same host. Though clinical studies suggest that infection by more than one unrelated respiratory virus may alter disease severity, animal models in which we can control the doses, timing, and strains of coinfecting viruses are critical to understand how coinfection affects disease severity. In this study, we compared gene expression and immune cell recruitment between two pairs of viruses (RV/PR8 and RV/PVM) inoculated sequentially in mice that both result in reduced severity compared to lethal infection by PR8 or PVM alone. Reduced disease severity was associated with suppression of inflammatory responses in the lungs. However, differences in disease kinetics and host and viral gene expression suggest that protection by coinfection with RV may be due to distinct molecular mechanisms. Indeed, we found that antiviral cytokine signaling was required for RV-mediated protection against lethal infection by PR8, but not PVM.

## Introduction

The detection of more than one virus in respiratory samples is quite common, especially among pediatric patients (1–4). There are differences in the outcomes of coinfection - whether it results in increased, decreased, or no effect on disease severity - that likely reflect different virus parings, patient populations, and study criteria. For example, coinfection with influenza B virus was found to increase the severity of seasonal influenza A virus, while other virus pairings did not reach statistical significance (2). Another study found increased rates of hospitalization, but not other measures of clinical severity, associated with viral coinfections (1). In contrast, Martin et al. found that patients with one virus detected had increased risk of severe disease than those with multiple viruses detected, and some virus pairings were associated with lower viral loads in coinfected patients (5). Coinfection by non-SARS-CoV-2 viruses has been detected in COVID-19 patients, but the impact on disease severity is not well-understood (6–8). Recently, RV has been shown to inhibit replication of SARS-CoV-2 in cultured human airway epithelial cells (9). Despite differences among studies, it is clear that coinfecting viruses have the potential to alter pathogenesis and disease outcomes. While clinical studies illuminate this potential, model systems in which the virus pairs, doses, and timing of coinfection can be controlled are critical to understand how coinfection alters pathogenesis in the respiratory tract. Therefore, we developed a mouse model using pairwise combinations of respiratory viruses from different families for this purpose (10).

We previously found that inoculation of mice with a mild respiratory virus (rhinovirus strain 1B [RV] or mouse hepatitis virus strain 1 [MHV-1]) attenuates the severity of a subsequent infection by influenza A virus (strain PR8) (10). Although pre-infection does not prevent PR8 replication, it leads to faster viral clearance and resolution of pulmonary inflammation. Protection from severe disease by viral coinfection has also been demonstrated by other groups. Similar to our study, Hua et al. found that a nasal-restricted infection by MHV-1 protects mice from lethal infection by PR8 and mouse-adapted severe acute respiratory syndrome coronavirus (SARS-CoV) (11). Furthermore, they found that protection is associated with enhanced macrophage recruitment to the lungs and up-regulation of SARS-CoV-specific CD4+ and CD8+ T cell responses (11). Inhibition or delay of viral shedding also occurs in ferrets during sequential inoculations with antigenically similar or dissimilar strains of influenza A and B viruses (12). A 2009 pandemic influenza A (A(H1N1)pdm09) virus prevents subsequent infection of ferrets by human respiratory syncytial virus (RSV) when the viruses are given within a short time frame (13). In contrast, RSV does not prevent replication of A(H1N1)pdm09 in ferrets, but reduces morbidity as determined by weight loss (13). Inhibition of RSV by prior inoculation with influenza A virus has also been demonstrated in mice (14).

In this study, we aimed to evaluate whether RV-mediated disease attenuation was specific to PR8, or generalizable to other respiratory viral infections. We found that RV reduced the severity of an additional lethal respiratory virus, pneumonia virus of mice (PVM). PVM is a relative of human RSV in the family *Pneumoviridae*, genus *Orthopneumovirus*. Infection of mice with PVM results in severe disease that shares clinical features of the most severe infections by RSV, including infection of the bronchiolar epithelium, granulocytic inflammation, and pulmonary edema (15–17). Lethality upon infection of mice with PR8 or PVM is mediated largely by dysregulated inflammatory responses, rather than overt damage due to viral replication (18–21). Interestingly, there are differences in the types of immune responses that mediate protection against these two viruses. Type I and III interferon (IFN) signaling and alveolar macrophages are protective in PR8-, but not PVM-, infected mice (18, 20, 22–25). Conversely, plasmacytoid dendritic cells are required for protection in PVM-, but not PR8-, infected mice (26, 27). Interleukin-6 (IL-6) limits the severity of influenza A virus infection in mice, while it exacerbates disease in PVM-infected mice (28–30). Multiple mechanisms reduce the severity of both PR8 and PVM infections, including mucin production and inhibition of inflammatory responses (31–37). Based on these complex differences in immunity to infection with PR8 and PVM, we used a global gene expression approach combined with flow cytometry to evaluate potential mechanisms whereby pre-inoculation with RV reduces the severity of PR8 and PVM.

## Results

### Inoculation of mice with RV reduces the severity of PVM infection

We previously showed that inoculation of mice with RV completely prevented mortality and reduced morbidity of a lethal infection by influenza A virus, strain PR8 (10). Protection against lethal PR8 infection was most effective when RV was given two days before PR8, but significant disease attenuation was also seen when mice were inoculated with RV and PR8 concurrently. To determine if RV would attenuate disease by a second lethal respiratory viral pathogen, we inoculated mice with media (mock) or RV two days before, simultaneously with, or two days after inoculation with PVM. Mice were monitored daily for mortality, weight loss, and clinical signs of disease for 14 days after PVM inoculation.

Infection with PVM alone (mock/PVM) was 100% lethal with all mice succumbing to infection or reaching humane endpoints by day 8 (Fig 1A). PVM-infected mice had rapid weight loss (Fig 1B) and exhibited clinical signs of disease, including ruffled fur, hunched posture, labored breathing, and lethargy. Mice that were inoculated with RV two days before (RV/PVM) or simultaneously with (RV+PVM) PVM were completely protected from mortality (Fig 1A), weight loss (Fig 1B), and clinical signs of disease. However, mice that received RV two days after PVM (PVM/RV) had equivalent disease severity as mock/PVM-infected mice. Although inoculation with RV reduced the severity of both PR8 (10) and PVM (Fig 1), protection was more effective against PVM. Importantly, complete protection against morbidity in addition to mortality was seen when RV was given two days before or concurrent with PVM (Fig 1). In contrast, RV prevented mortality, but not morbidity, when given two days before PR8 and inoculation of RV and PR8 concurrently was less effective at reducing disease severity (10).

**Fig 1.**
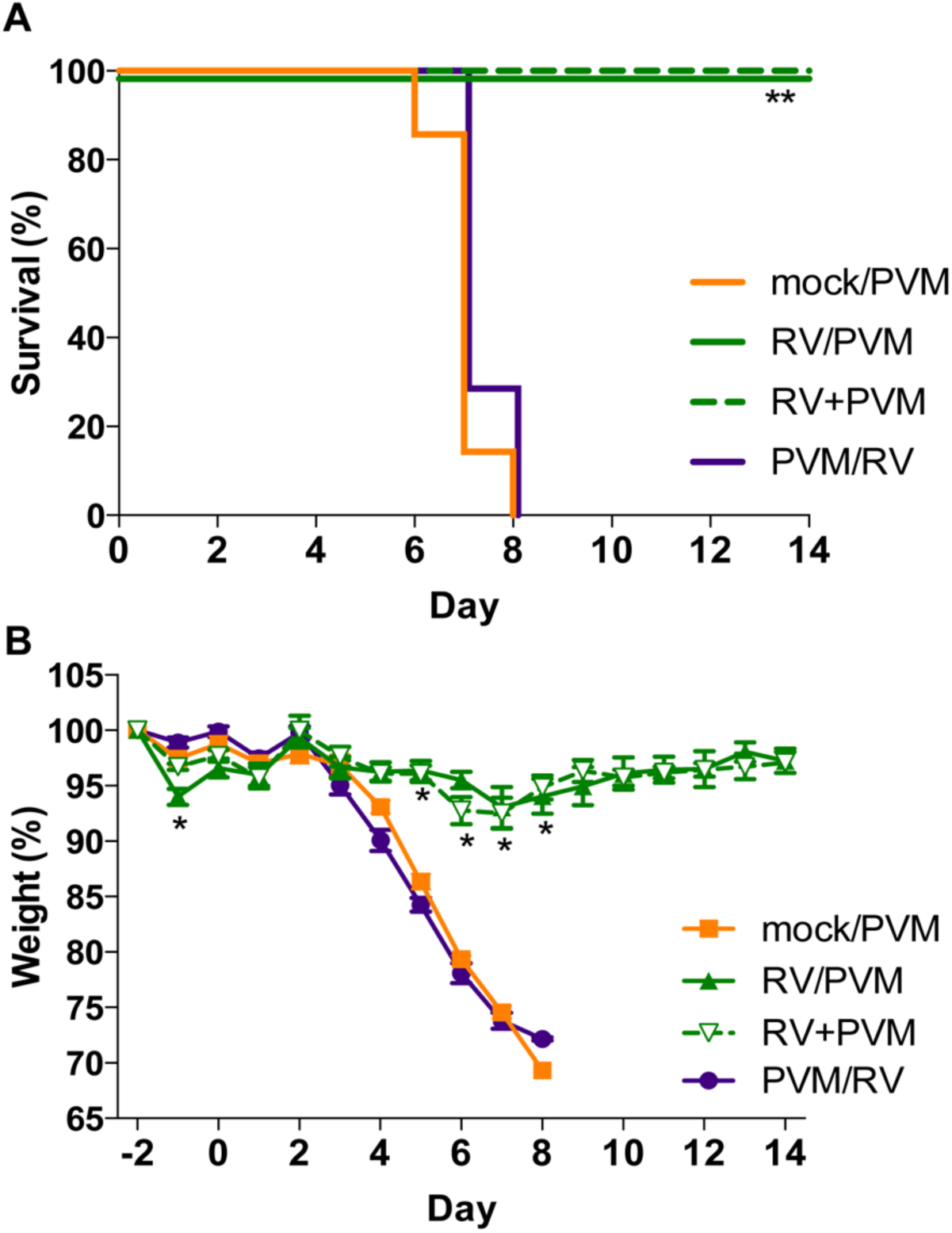
Inoculation with RV reduces the severity of PVM infection. Groups of 7 BALB/c mice were either mock-inoculated (mock/PVM) or inoculated intranasally with 7.6 x 10^6^ TCID_50_ units of RV two days before (RV/PVM), simultaneously with (RV+PVM), or two days after (PVM/RV) 1.0×10^4^ TCID_50_ units of PVM. Mice were monitored daily for (A) mortality and (B) weight loss. Data are representative of two independent experiments. Statistical significance compared to mock/PVM was determined by (A) Log-rank Mantel-Cox test and (B) student’s t-test corrected for multiple comparisons using the Holm-Sidak method. P-value *p≤0.05, **p≤0.005, ***p≤0.001.

### Infection by PR8 and PVM induce different gene expression signatures in mouse lungs over time

To determine potential mechanisms of protection mediated by RV against PR8 and PVM, we undertook a comprehensive transcriptome analysis of mouse lungs (Fig 2). Mice were inoculated with RV two days before PR8 or PVM and RNA isolated from the right lobes was analyzed on days 0, 2, 4, and 6 after PR8 or PVM inoculation. Single virus-infected mice were mock-inoculated two days before PR8 or PVM. Weight loss was monitored daily to test for consistency with our previous coinfection studies. RV-mediated protection against PR8 was not evident by 6 days post-infection (S1 Fig) though both mock/PR8 and RV/PR8 groups experienced weight loss at a rate similar to our previous study (10). In contrast, complete protection against weight loss was evident in RV/PVM treated mice (S1 Fig).

**Fig 2.**
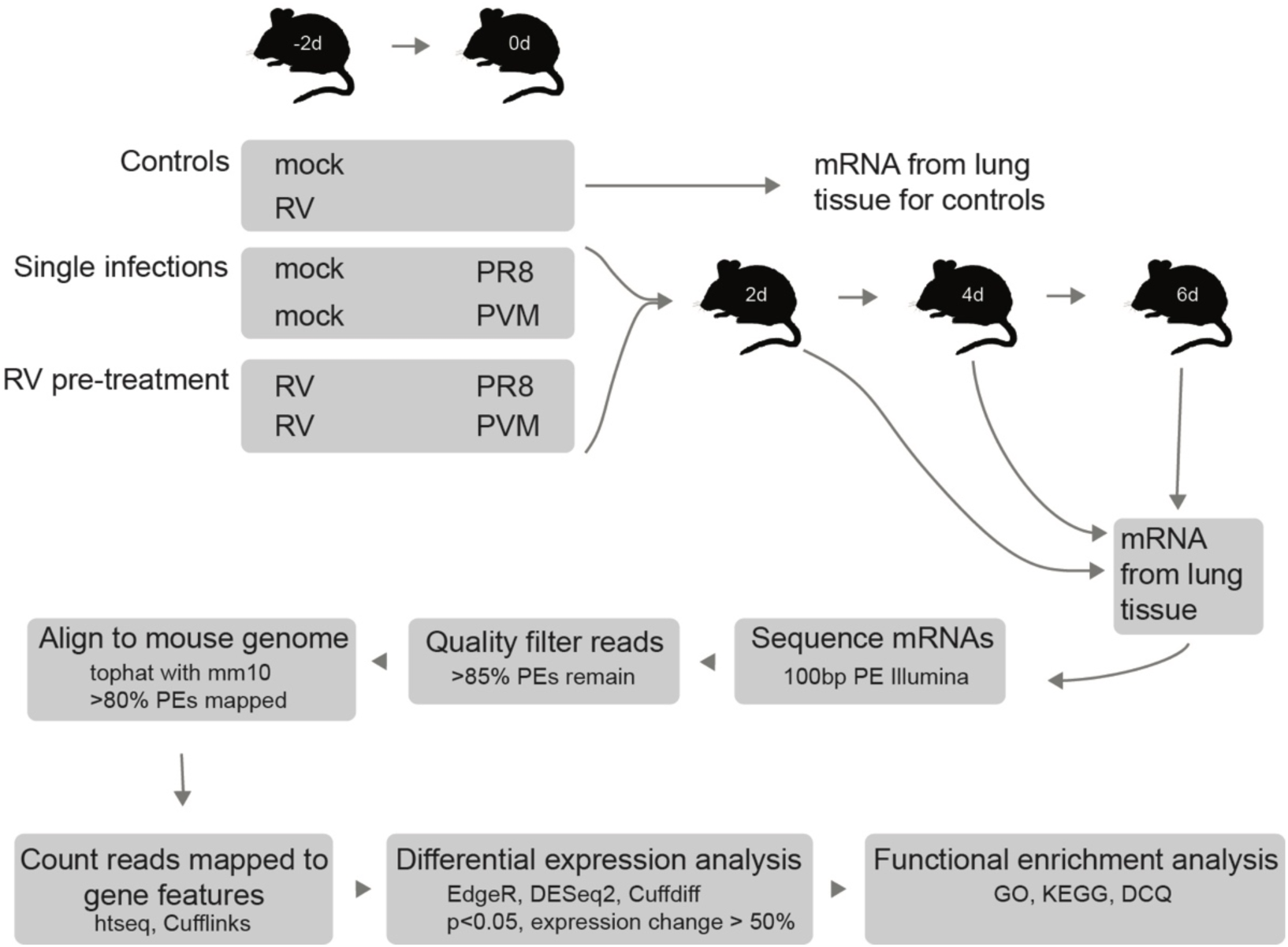
Experimental outline of transcriptome study. Mice were inoculated with RV or saline (mock) two days before challenge with PR8 or PVM. RNA was extracted from lung tissue from three mice per group on day 0 for mock and RV treated controls and days 2, 4, and 6 for mice receiving PR8 or PVM on day 0 and analyzed by RNA-seq.

Lung mRNA from 3 mice per infection group and time point was collected and sequenced on the Illumina platform. Of the 14.6 to 55.3 (36.4 mean) million reads obtained for each sample, 83-91.2% were uniquely mapped to the mm10 mouse genome (GCA_000001635.2). The expression of 24,243 and 24,421 genes were measured using Cufflinks and HTSeq, respectively. We conservatively included differentially expressed genes (DEGs) by requiring that they be identified by Cuffdiff, EdgeR, and Deseq2 as having a log_2_ fold change value of at least 0.5 and *p*-value less than 0.05.

A principal component analysis (PCA) was done with the 500 most highly variable genes to visualize variation across infection groups, time points, and replicates. For the most part, replicate samples clustered together (Fig 3A). Most variation in host gene expression was explained by the time elapsed since viral infection (Fig 3A). However, differences between viruses were also apparent. Infection by PVM caused fewer gene expression changes at early time points than PR8-infected mice, as seen by mock/PVM and RV/PVM day 2 samples clustering closely with mock and RV day 0 samples, respectively (Fig 3A, blue and purple points). By day 4, the PVM-infected mice had very different gene expression profiles than mock-inoculated mice on day 0. This delayed response to PVM infection is observed in other studies (18, 38). In contrast, PR8 infection dramatically altered gene expression in mice by day 2 (Fig 3A, light green points). Early changes in host gene expression induced by PR8, but not PVM, corresponded with earlier detection of PR8-specific transcripts (Fig 4). Moreover, the RV/PVM and mock/PVM samples were very different from each other by day 4 (Fig 3A, brown points). RV/PR8 and mock/PR8 groups had similar gene expression signatures until day 6 (Fig 3A, orange points). This corresponds with the similar disease kinetics in the mock/PR8 and RV/PR8 groups early during infection (10) (S1 Fig).

**Fig 3.**
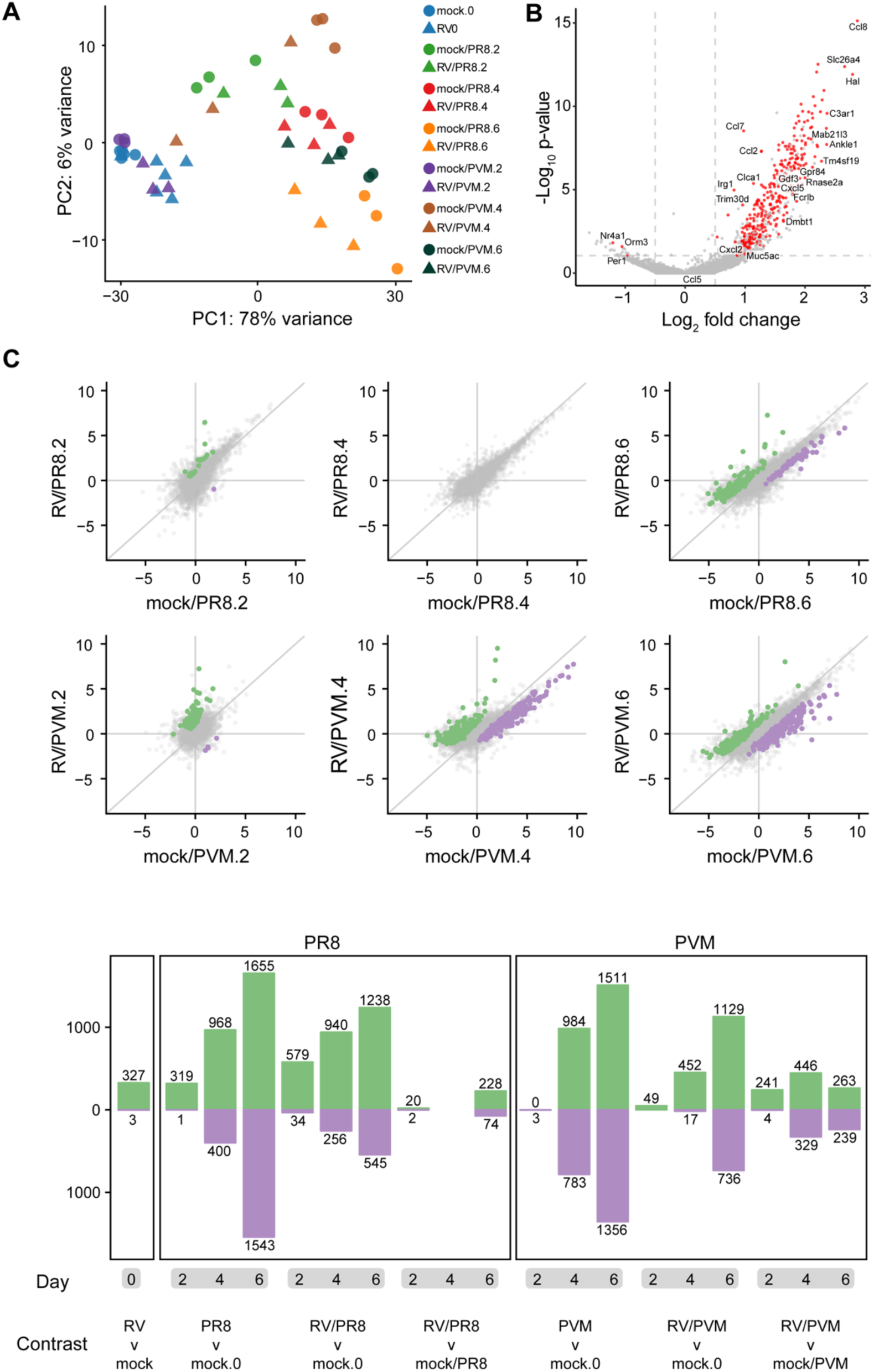
Patterns of mouse gene expression in lungs upon infection with and without RV pre-treatment. (A) Principal component analysis of RNA-seq data shows that the greatest variance in gene expression is primarily due to time since infection. The largest distances in single virus-infected vs. RV-treated groups are mock/PVM vs. RV/PVM on day 4 and mock/PR8 vs. RV/PR8 on day 6. (B) Volcano plot showing gene expression changes between mock-inoculated and RV-inoculated mice 2 days post-infection. Only DEGs identified by edgeR, DESeq2, and Cufflinks were considered significant (red points). (C) The scatter plots show the log2 fold change values of all genes compared to mock-inoculated mice on day 0. Genes upregulated in RV-treated compared to the single-virus infected mice are colored green. Down-regulated genes are colored purple. (D) Numbers of DEGs in all pairwise comparisons. The number of genes upregulated in the infected (versus mock) or RV-treated (versus mock or single) mice are shown in green. Downregulated genes are shown in purple. See Supplemental Table 1 for lists of gene names and log2 fold change values.

**Fig 4.**
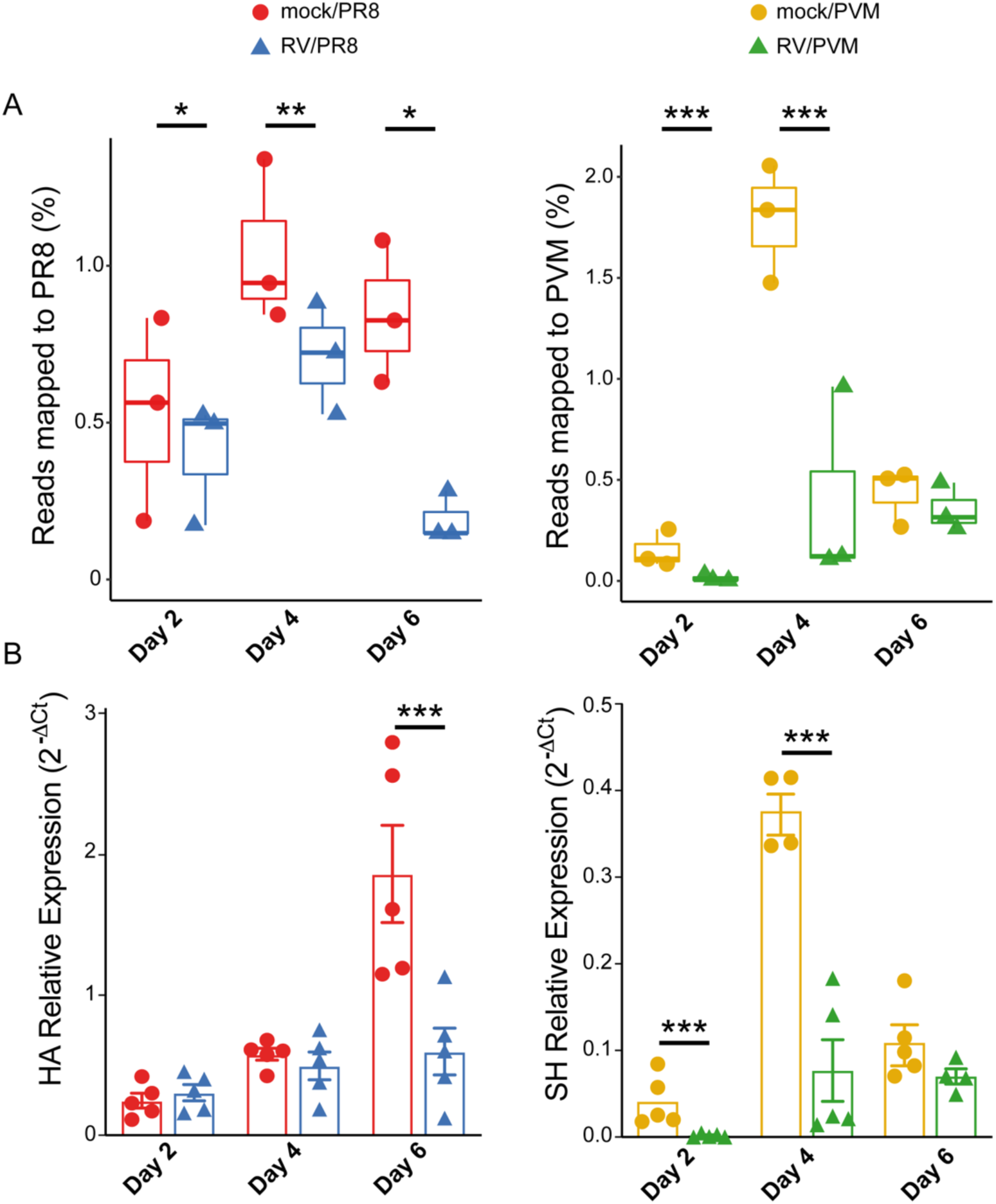
Viral gene expression in single virus-infected and RV-treated mouse lungs. (A) RNA-seq quantification of viral mRNAs. For each group, the three replicates are shown with box plots indicating the quantile values of percent non-mouse reads mapped to viral genomes. Asterisks indicate significantly different treatment pairs as determined by a generalized linear model, see Supplemental Table 3. P-value *p<0.05, **p<0.01, ***p≤0.001. (B) RT-qPCR analysis of viral RNAs. Replicates from individual mice are shown with the mean and standard error indicated and is representative of at least two replicate assays. Statistical significance between single virus-infected and RV-treated mice at each time point was determined using unpaired t-tests corrected for multiple comparisons using the Holm-Sidak method. ***p≤0.001.

We next calculated differential gene expression levels in samples from RV-inoculated mice compared to mock-inoculated mice on day 0 (Fig 3B). Similar to the PCA, gene expression was significantly changed 2 days post-infection with RV and the response was markedly skewed towards up-regulation. DEGs identified in all infection groups vs. mock were compared between single virus-infected and RV-treated mice at each time point (Fig 3C). Gene expression changed over time, as samples taken 2 days post-infection were most similar to mock-inoculated controls and samples taken 6 days post-infection were most different (Fig 3C, 3D). The transition of point clustering in Fig 3C from around the origin at day 2 to spread along the 1:1 line on day 6 illustrates this delayed gene expression response to viral infection. Genes that were differentially expressed between single virus-infected and RV-treated mice also displayed time-dependent clustering shifts in both the PVM and PR8 experiments (Fig 3C). At 2 days post-infection, the expression values of DEGs indicated that the samples from RV/PR8 and RV/PVM groups were more different from mock than the samples from single virus-infected mice (points are spread along the y-axis). At 6 days post-infection, DEGs differed more between mock-inoculated and single virus-infected mice than between mock and RV-treated mice (points spread along x-axis). This dampening of DEGs in RV/PR8 and RV/PVM groups suggests that infection by a lethal virus is ablated not only in virulence, but also in host gene expression response.

A summary of the numbers of DEGs across comparisons of all virus-infected to mock-inoculated mice and between single virus-infected vs. RV/PR8 or RV/PVM treated mice is shown in Fig 3D and lists of these DEGs are included in Supplemental Table 1. Again, the numbers of DEGs reflect the delayed response to PVM infection, dampened gene expression in RV/PR8 and RV/PVM groups of mice, and relative similarity in gene expression in mock/PR8 and RV/PR8 infected mice early in infection.

### Viral gene expression

Sequencing reads that did not map to the mouse genome were aligned with viral genomes. An insignificant number of reads mapped to the RV genome. This was expected because the abundance of positive-sense RV RNA and viral titers peak 24 hours after inoculation of BALB/c mice (39). The earliest time we analyzed was 48 hours (day 0) after inoculation, at which point RV RNA would likely represent a very low proportion of the total RNA. We confirmed this by quantifying infectious RV in homogenized lung tissue and broncho-alveolar lavage fluid (BALF) 24 h and 48 h after RV inoculation, days -1 and 0, respectively. In agreement with published studies and our RNA-seq results, low levels of RV were detected on day -1, but not day 0 (S10 Fig). Thus, we cannot conclude that RV is replicating in mouse lungs despite having a significant impact on host gene expression.

Many reads mapped to PR8 and PVM genomes from the mice infected with these viruses (Fig 4). Regression analyses were performed to identify significant changes over time (S3 Table). Pre-treatment with RV did not prevent PR8-specific gene expression, but reads mapped to PR8 were lower in RV/PR8 mice at all time points (Fig 4A). When this analysis was expanded to additional animals assayed by RT-qPCR for the HA gene, RV/PR8 infected mice only had significantly lower viral gene expression compared to mock/PR8 infected mice on day 6 (Fig 4B), which was also the time point that was most dramatically different in RNA-seq reads. Similarly, we previously showed that infectious PR8 titers in the lungs were equivalent in mock/PR8 and RV/PR8 infected mice on days 2 and 4 after PR8 inoculation (10). However, incoulation with RV led to earlier clearance of PR8 (by day 7), corresponding with the dramatically lower PR8 mRNAs seen on day 6 in the present study. These findings confirm that infection with RV does not prevent subsequent infection by PR8, rather it reduces viral gene expression, specifically late during infection. Our sequencing protocol only captured polyadenylated RNAs, thus the PR8 reads represent viral mRNAs and not complementary (cRNA) or genomic (vRNA) viral RNAs, while RT-qPCR was performed on total RNA. Based on individual gene mapping, the PR8 reads predominantly mapped to the nucleoprotein (NP) and hemagglutinin (HA) mRNAs (S2 Fig), which are known to be expressed at high levels during infection (40–42).

PVM gene expression was highest on day 4, but was robustly suppressed in RV-inoculated mice (Fig 4). This trend was confirmed by RT-qPCR quantification of the PVM SH gene (Fig 4B) and quantification of infectious virus (see Fig 8E), below. This suggests that RV may limit PVM infection early, which corresponds with the more effective prevention of weight loss in RV/PVM (Fig 1) compared to RV/PR8 (10) infected mice (S1 Fig). All viral genes were detected in mice infected by PVM alone (mock/PVM) on day 4, with the genes that express the attachment (G), nucleoprotein (N), non-structural 2 (NS2), fusion (F), phosphoprotein (P), and matrix (M) proteins at the highest levels (S2 Fig). This does not strictly follow the gradient of mRNA levels corresponding with gene order that is expected from pneumoviruses (43), which likely reflects post-transcriptional differences in mRNA stability and the heterogeneous nature of collecting cells at different stages of the virus replication cycle.

### RV induces innate immune response prior to secondary viral infection

We compared gene expression in lung tissue of mock-inoculated mice to mice inoculated with RV for 2 days (RV, day 0). Of the 24,421 genes compared, we identified 330 DEGs, of which only three were down-regulated (Fig 3B, 3D, and S1 Table). To get a functional picture of the RV-treated lung on day 0, we identified enriched GO terms and KEGG pathways in the 330 DEGs. The most highly enriched terms on this list suggest that changes in the regulation of cell cycle or cell division were occurring in RV-compared to mock-inoculated mice; nearly all of the top 50 most enriched terms involved chromosome remodeling and mitosis (S2 Table), suggesting RV-induced increase in cellular proliferation. This list of enriched terms was quite different than the processes that were differentially regulated in PR8-or PVM-infected mice. Up-regulation of cell division-associated genes could be occurring in epithelial cells in conjunction with repair, immune cells recruited to the lungs, or both. In addition to cell division, multiple immune response-related GO terms related to type I IFN signaling, chemokine signaling, and immune cell chemotaxis were enriched in RV-treated mice (S2 Table). RV induced up-regulation of several chemokine genes, including those that recruit monocytes, neutrophils, NK cells, T cells, B cells, and eosinophils (Ccl-2, 3, 6, 7, 8, 12, 17, 22, Cxcl-1, 3, 5, 9, 10, 13).

### Host gene expression changes in RV/PR8 and RV/PVM infected mice

We next identified DEGs specific to mice exposed to two viruses (RV/PR8 or RV/PVM) vs. mock-inoculated mice that were not differentially expressed in single virus-infected (mock/PR8 or mock/PVM) vs. mock-inoculated mice at the same time points. The expression data for these three gene sets (unique to RV/PR8, unique to RV/PVM, shared by RV/PR8 and RV/PVM) are provided in supplemental figures (Fig S3-S5). Genes with shared up-regulation in RV-inoculated animals, regardless of time point and second virus, included those in the mucin biosynthesis pathway, MHC class II genes, and immunomodulatory genes. Genes with increased up-regulation in mice infected with RV/PVM compared to mock/PVM early (days 2 and 4) included Mgl2, Ccl6, Agr2, H2-M2, and Muc5b. Genes that had higher up-regulation in the single virus infections were predominantly increased late (days 4 and/or 6) and included genes associated with inflammation (Angptl4) or pulmonary fibrosis (Fosl2, Pappa, Sphk1). These genes were common to both PR8 and PVM infections and likely result from excessive inflammation and tissue damage within the infected lungs. Additional genes associated with inflammation and fibrosis (Nfkbia and Mmp9) or stress responses (Hspb8, Nupr1) were reduced in RV/PVM-compared to mock/PVM-infected mice. Finally, a set of genes were up-regulated in both mock/PVM-and RV/PVM-infected mice, but to a higher level in mock/PVM-infected mice (S100a9, Prss22, Fga, Krt17). These genes are also largely involved in inflammation and tissue damage and repair processes.

Genes involved in goblet cell metaplasia and mucin production were specifically increased in both RV/PVM and RV/PR8 infected mice compared to the single virus-infected mice at various time points. These included the major gel-forming airway mucins (Muc5ac and Muc5b), a disulfide isomerase (Agr2) required for mucin folding and polymerization, and a chloride channel regulator (Clca1) required for proper hydration of mucus. Additional ion channels (Slc6a20a), aquaporins (Aqp9), and mucus-associated proteins (Itln1) also had increased expression in lungs from RV-treated mice. In addition to the mucin-related genes shared with RV/PR8, RV/PVM infected mice had increased expression of a transcription factor (FoxA3) that promotes goblet cell metaplasia and mucus production. Muc5ac and Clca1 had increased expression in RV-treated mice at all time points (Fig 5).

**Fig 5.**
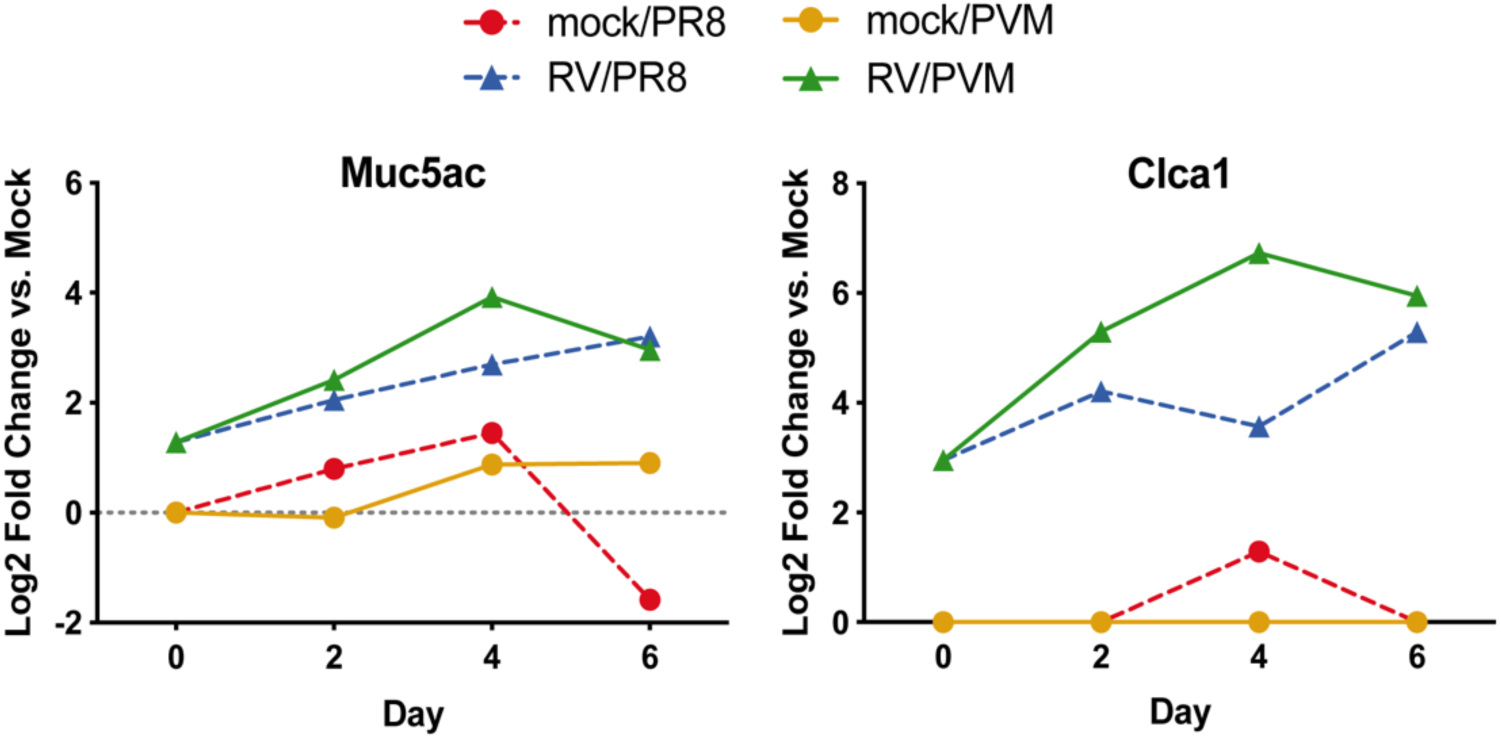
Mucin-related genes have increased expression in RV-inoculated mice across all time points. Log2 fold change values compared to mock day 0 are shown for mucin 5ac (Muc5ac) and chloride channel accessory 1 (Clca1) genes, which are required for mucus production and hydration, respectively.

Based on the importance of type I IFN and inflammatory signaling pathways in the pathogenesis of viral infections, we generated heatmaps demonstrating relative gene expression across all groups for genes in the Hallmark Interferon Alpha Response and Inflammatory Response gene sets from MSigDB (44, 45). Overall, IFN-response genes had higher expression in RV/PR8 vs. mock/PR8 mice on day 2 and lower expression by day 6 (Fig 6A and S6 Fig). This pattern corresponds with our previous study that demonstrated increased expression of IFN-β only on day 2 in RV/PR8 compared to mock/PR8 infected mice (10) and also the lower PR8 gene expression in RV/PR8 infected mice (Fig 4). In contrast, mock/PVM and RV/PVM infected mice had delayed up-regulation of IFN-response genes and mock/PVM infected mice had dramatically higher expression of IFN-response genes compared to RV/PVM on day 4 (Fig 6B and S6 Fig). This pattern was validated by RT-qPCR analysis of IFN-β expression (Fig 8D) and corresponds with the overall delayed PVM-induced gene expression and reduced levels of PVM RNA in RV/PVM infected mice (Fig 4). Inoculation with RV induced expression of a small subset of IFN-response genes (Il7, Ifi27, Lamp3, Cd74, Ifi30, and Lpar6) on day 0, which was maintained in RV/PVM-infected mice on day 2 (Fig 6B). While more variation in gene expression patterns was observed in the inflammatory response gene set, a large number of genes followed the same trends seen for IFN-response genes, i.e., a largely dampened response in RV-treated mice especially at later time points (Fig 6C, 6D and S7 Fig). By day 6, mock/PR8-infected mice had strongly up-and down-regulated expression of inflammatory response genes, while RV/PR8-infected mice had muted changes in expression of these genes (Fig 6C). A large subset of inflammatory response genes had similar patterns as IFN-response genes in mock/PVM and RV/PVM mice, including delayed expression and a muted response in RV-treated mice (Fig 6D). This subset includes important inducers of inflammatory responses such as Nfkb1, Nfkbia, Rela, and Tlr3. Other subsets of genes (e.g., Ccl17 and Ccl22) had early up-regulation in RV-inoculated mice on day 0, which was maintained on day 2 after infection with PVM, and reduced in RV/PVM infected mice on days 4 and 6. In contrast with IFN-response genes, subsets of inflammatory response genes had higher expression in RV/PVM compared to mock/PVM infected mice throughout the course of infection. This suggests that, despite low virus levels (Fig 4) and no clinical signs of disease (Fig 1 and S1) and largely muted host gene expression, expression of some host genes is increased in RV/PVM-infected mice.

**Fig 6.**
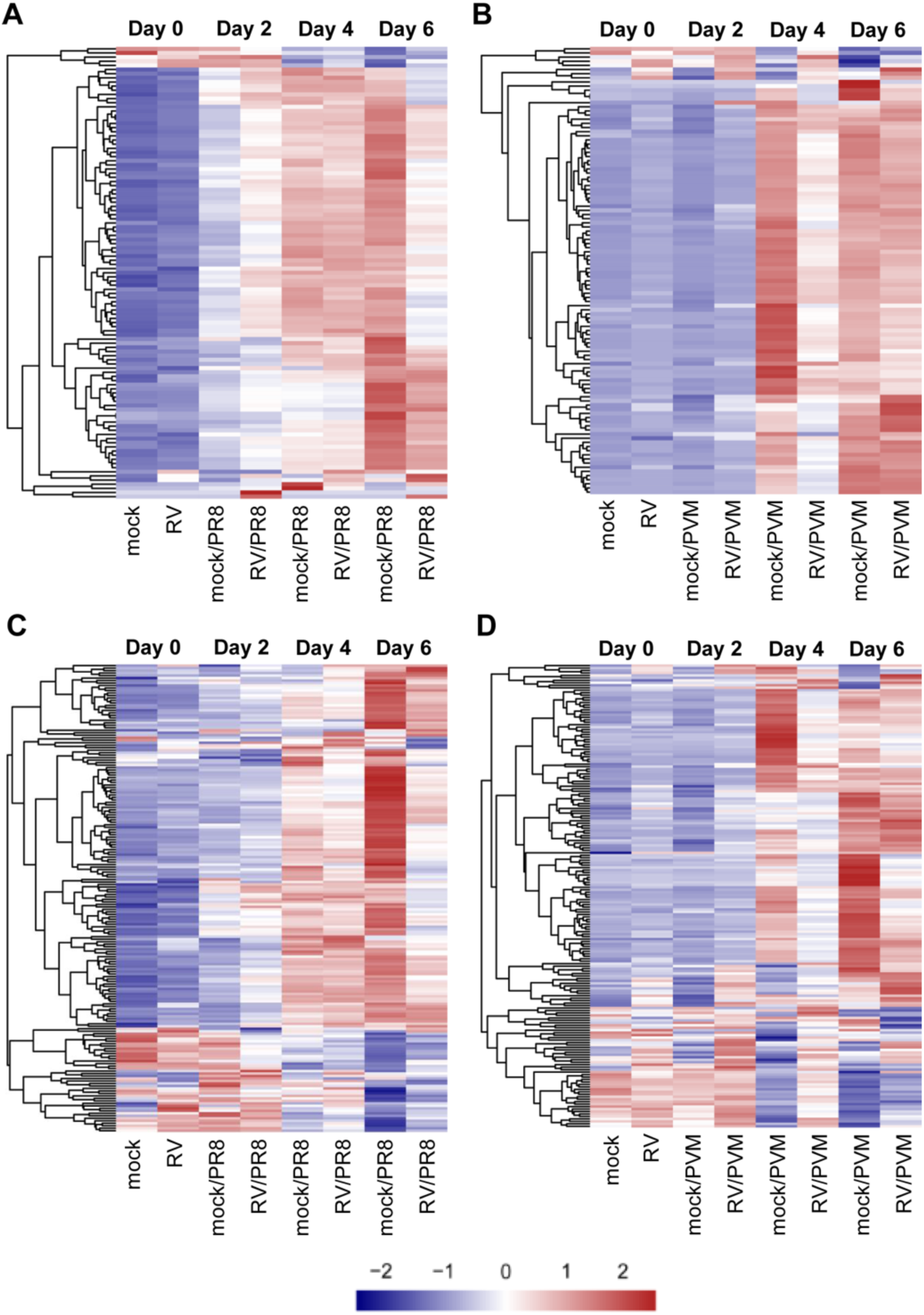
Expression of MSigDB Hallmark Interferon Alpha Response and Inflammatory Response gene sets. Heatmaps of Hallmark Interferon Alpha Response genes showing the relative expression (Z-scores) of genes for the (A) PR8 or (B) PVM infection RNA-seq time course. Heatmaps of Hallmark Inflammatory Response genes showing the relative expression (Z-scores) of genes for the (C) PR8 or (D) PVM infection RNA-seq time course. All heatmaps represent DESeq2 normalized counts where each row represents an individual gene. The genes were ordered by hierarchical clustering, which is shown on the left side of each heatmap. The colors (blue < white < red) represent the Z-score and a more intense color indicates a lower (blue) or higher (red) relative expression of that gene in that condition. Fully labeled heatmaps can be found in S6 and S7 Fig.

### Flow cytometry analysis

Immune cells recruited to the lungs are affected by and contribute to the gene expression signatures seen in whole lung tissues. To analyze differences in immune cell recruitment to the lungs PR8 and PVM infected mice with and without RV pre-treatment, we quantified innate immune cells in the left lobes by flow cytometry at the same time points as our gene expression analyses (Fig 7) and performed regression analyses to identify significant changes over time (S3 Table). Total cell counts from mock/PVM-and RV/PVM-infected mice were fairly consistent across all time points, while total cells in mock/PR8 and RV/PR8 infected mice increased by day 6 (S8 Fig). CD11b+ was used to differentiate leukocytes from resident lung cell populations. CD11b+ cells accounted for the increase in mock/PR8 and RV/PR8 infected lungs (Fig 7A). In contrast, while CD11b+ cells increased in the lungs of mice infected with mock/PVM, RV/PVM-infected mice had reduced recruitment of these cells on days 4 and 6 (Fig 7E). Neutrophils were gated based on high expression of CD11b and Ly6G and the remaining cells were identified as alveolar macrophages (CD64+/SiglecF+) and interstitial macrophages (CD11b+/CD64+/SiglecF-) (S8 Fig) (46, 47).

**Fig 7.**
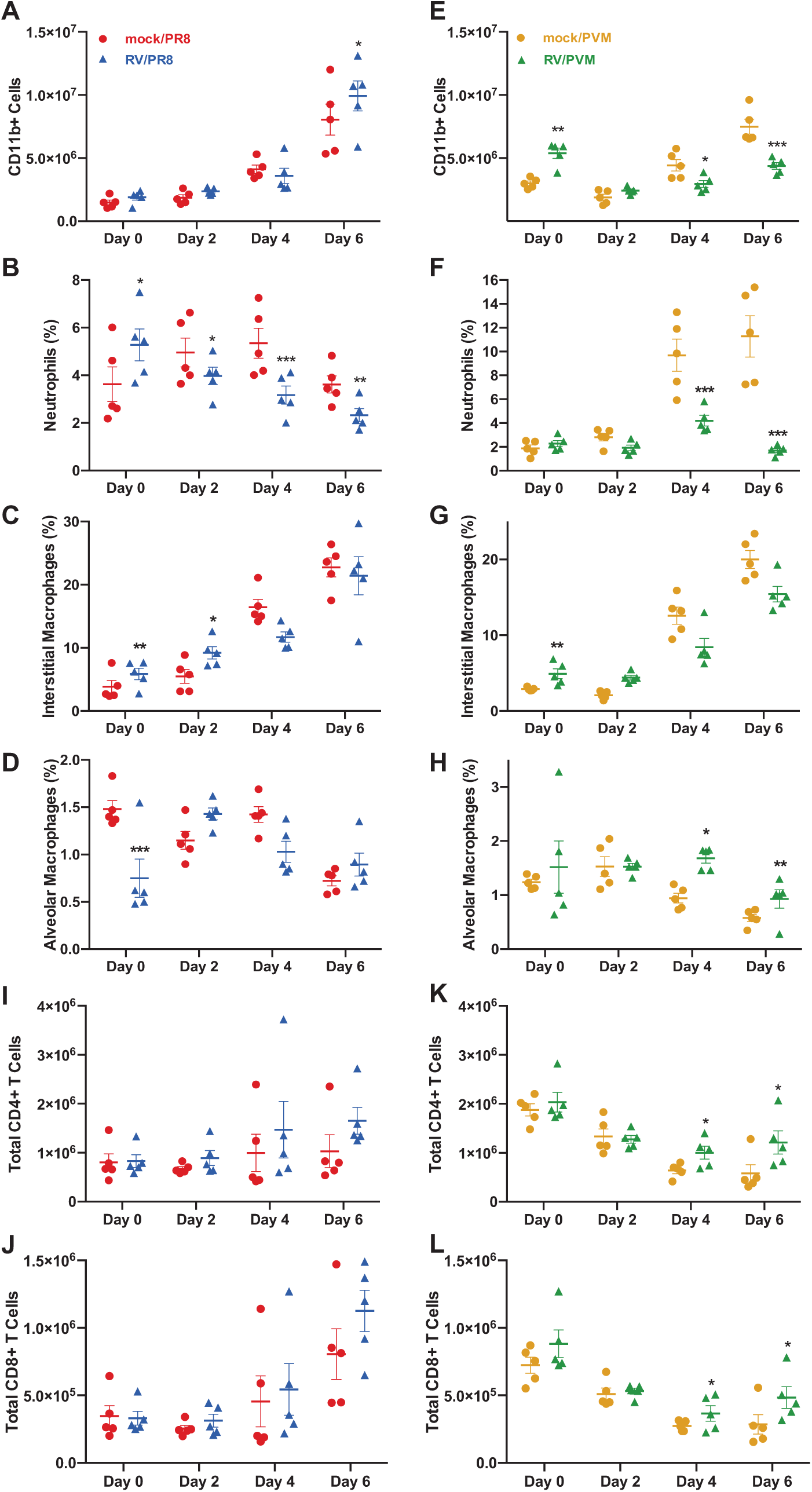
Flow cytometry analysis of innate immune cell populations in the lungs upon viral infection and RV-treatment. Mice were inoculated with mock or RV on day -2 and PR8 (A-D, I, J) or PVM (E-H, K, L) on day 0, followed by analysis of specific cell populations in lung homogenates by flow cytometry, including total CD11b+ cell counts (A, E), percentages of CD11b^hi^/Ly6G^hi^ neutrophils (B, F), CD11b+/CD64+/SiglecF-/Ly6G-interstitial macrophages (C, B), and CD64+/SiglecF+/Ly6G-alveolar macrophages (D, H), and total numbers of CD3+/CD4+ (I, K) and CD3+/CD8+ (J, L) T cells. Asterisks indicate significantly different treatment pairs, see Supplemental Table 3. P-value *p<0.05, **p<0.01, ***p≤0.001.

The proportions of neutrophils and interstitial macrophages followed the same trends as total CD11b+ cells in PVM-infected mice, with lower proportions of these cells on days 4 and 6 in RV/PVM-infected mice (Fig 7F, 7G). In contrast, interstitial macrophages were increased in RV/PVM, compared to mock/PVM-infected mice, early in infection. This indicates that pre-treatment with RV stimulates early recruitment of CD11b+ cells, specifically interstitial macrophages, while limiting recruitment of inflammatory cells later in infection. PR8-infected mice had similar trends, however the differences between mock/PR8 and RV/PR8 groups were less dramatic (Fig 7B, 7C). Neutrophil numbers were suppressed in RV/PR8-infected mice compared to mock/PR8-infected mice through-out the time course (Fig 7B). The interstitial macrophage proportions in mock/PR8- and RV/PR8-infected mice increased over time similarly to the total CD11b+ populations (Fig 7C). The lower proportions of neutrophils and interstitial macrophages at later time points in RV-treated mice corresponded with mRNA levels for chemokines. This was predominantly the case for neutrophil chemokines Cxcl1 and Cxcl2 and macrophage chemokines Ccl2 and Ccl7 (S9 Fig). These chemokines were generally lower in RV/PR8-infected mice on day 6 and RV/PVM-infected mice on days 4 and 6 compared to mock/PR8- and mock/PVM-infected mice, respectively.

There were no clear trends in alveolar macrophage numbers in mock/PR8- and RV/PR8-infected mice (Fig 7D), though their proportions were significantly higher in RV/PVM-infected mice compared to mock/PVM-infected mice on days 4 and 6 (Fig 7H). This is likely due to depletion of alveolar macrophages by PVM infection of these cells (48). A separate flow cytometry antibody panel was used to quantify CD4+ and CD8+ T cells in the lungs. T cell numbers in mock/PR8 and RV/PR8 infected mice increased over time, but the differences between the groups were not significant (Fig 7I, 7J). RV/PVM-infected mice had modest, yet significantly higher numbers of CD4+ and CD8+ T cells compared to mock/PVM-infected mice on day 6 (Fig 7K, 7L).

### IFNAR signaling is required for RV-mediated protection against PR8, but not PVM

To determine whether signaling by type I IFNs is required for RV-mediated reduction in disease severity, we treated mice with IFN-*αβ* receptor (IFNAR-1) blocking antibody on days -2 (with RV inoculation) and 0 (with PR8 or PVM inoculation). Mock/PR8-infected mice had similar weight loss and mortality with anti-IFNAR and control antibody treatments (Fig 8A). In contrast, treatment with anti-IFNAR antibody completely abrogated the dampening effects of RV on the morbidity and mortality of PR8 (Fig 8A). Thus, IFNAR signaling is required for the reduced disease severity observed in RV/PR8-infected mice. Mice that were inoculated with RV alone did not experience weight loss, clinical signs of infection, or mortality over 14 days when treated with either anti-IFNAR or control antibodies (Fig S10). Interestingly, there were no significant differences in weight loss and mortality between anti-IFNAR- and control antibody-treated RV/PVM-infected mice (Fig 8B). Thus, type I IFN signaling is not required for RV-mediated protection against lethal PVM infection.

**Fig 8.**
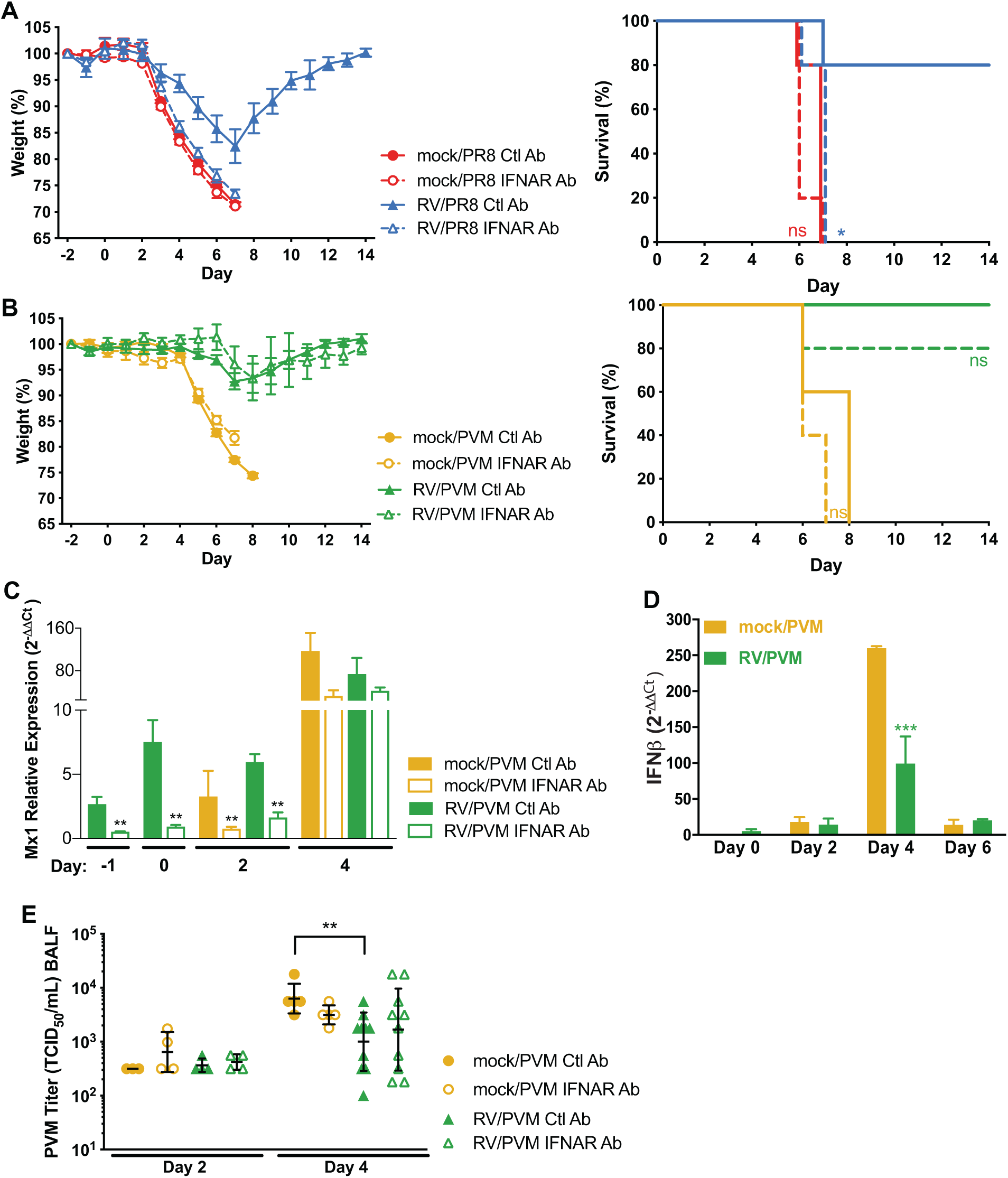
IFNAR signaling is required for RV-mediated protection in RV/PR8, but not RV/PVM, infected mice. Mice were treated with anti-IFNAR or an isotype control (Ctl) antibody (Ab) intranasally with viral inocula (mock or RV on day -2 and PR8 (A) or PVM (B-E) on day 0). Animal weights and survival were monitored in five mice per group for 14 days after inoculation with (A) PR8 or (B) PVM. Significant differences in survival between IFNAR Ab-vs. Ctl Ab-treated mice were determined using survival curve analysis by log-rank Mantel-Cox test. *p<0.05, ns=not significant, p>0.05. (C) Expression of IFN-induced gene, Mx1, was monitored to evaluate the effectiveness of IFNAR Ab treatment in mock/PVM and RV/PVM infected mice. Data shown are means and standard error of 4-5 mice per group and are representative of two replicate assays. (D) IFN-β RNA was quantified by RT-qPCR in 5 animals per group in mock/PVM and RV/PVM infected mice without antibody treatment. Data shown are means with standard error and are representative of two assays. (E) PVM titers in BALF were quantified by TCID_50_ assays. Data shown are from individual animals with the geometric means and standard deviations indicated. Statistical significance between groups within each time point was determined by unpaired t-tests corrected for multiple comparison by the Holm-Sidak method. **p<0.01, ***p<0.001.

To verify that IFNAR signaling was effectively inhibited in anti-IFNAR treated mice, we analyzed expression of an IFN-induced gene, Mx1, in mock/PVM and RV/PVM infected mice treated with anti-IFNAR or control antibody. Significant inhibition of Mx1 expression was seen through the course of anti-IFNAR treatment (days -1 through 2) with recovery of IFNAR signaling on day 4 after PVM infection (Fig 8C). In agreement with the weight loss and mortality data, inhibition of IFNAR signaling also had no significant effect on the viral load of PVM in mock/PVM or RV/PVM infected mice (Fig 8E).

### IFN-dependent protection in RV/PR8 infected mice is associated with reduced viral spread, neutrophilic inflammation, and histopathology in the lungs

Type I IFN signaling stimulates multiple downstream responses, including expression of antiviral genes, activation of innate and adaptive responses, and modulation of inflammatory responses. To better understand the role of IFNAR signaling in limiting disease severity in RV/PR8 infected mice, we analyzed PR8 replication, immune cell recruitment, and histopathology in RV/PR8-infected mice treated with anti-IFNAR or control antibodies. Anti-IFNAR treated mice had higher levels of PR8 in the lungs on days 2 and 6 than control antibody-treated mice, though the differences were not statistically significant (Fig 9A). Similarly, PR8 antigen was more widespread in the lungs of anti-IFNAR treated mice on days 2 and 6 post-infection (Fig 9B). However, IFNAR signaling did not completely prevent PR8 replication in RV/PR8-infected mice. This is in agreement with similar levels of PR8 RNA (Fig 4), viral loads, and antigen (10) in RV/PR8 infected mice compared to mock/PR8 infected mice. Inhibition of IFNAR signaling also resulted in higher titers of RV in BALF on day -1, with no infectious RV detected in IFNAR inhibited or control animals on day 0 (Fig S10).

**Fig 9.**
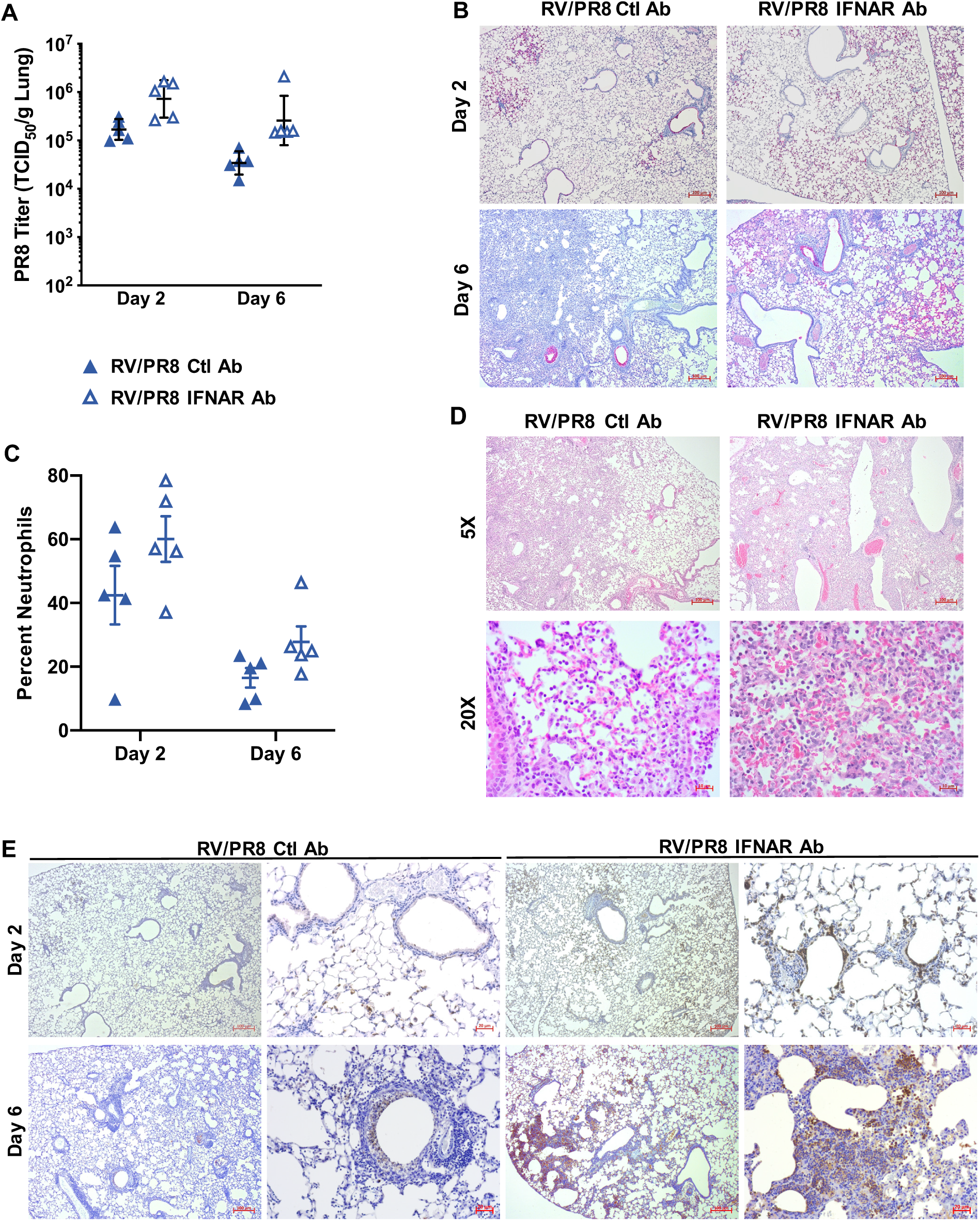
IFNAR-dependent protection in RV/PR8 infected mice is associated with reduced viral spread, neutrophilic inflammation, and histopathology in lungs. Mice were treated and infected as described in Fig. 8 legend. (A) PR8 was quantified in homogenized lungs by TCID_50_ assays. Data shown are from five individual mice per group with geometric mean and standard deviation indicated. Statistical significance between Ctl Ab and IFNAR Ab treated mice was determined using unpaired t-tests corrected for multiple comparisons using the Holm-Sidak method, and was not significant (p>0.05). (B) Viral antigen in lung tissues was visualized by IHC using antibody against PR8 HA protein followed by alkaline phosphatase with impact red substrate. (C) Cytospins from BALF were stained with HEMA 3 to identify airway neutrophils. At least 300 total cells were counted from duplicate cytospins and the percent neutrophils was calculated. Data shown are from individual mice with mean and standard error indicated. Statistical significance between Ctl Ab and IFNAR Ab treated mice was determined using unpaired t-tests corrected for multiple comparisons using the Holm-Sidak method, and was not significant (p>0.05). (D) Histopathology in lung tissues was visualized by hematoxylin & eosin staining on day 6 after PR8 infection. Arrows indicate examples of pulmonary hemorrhage. (E) Neutrophils in lung tissues were visualized by IHC using antibody against Ly6G followed by horseradish peroxidase and impact DAB substrate. Images in B, D, and E are representative of two mice per group and time point.

Mice treated with anti-IFNAR antibody had a slightly higher percentage of neutrophils in the airways compared to mice treated with a control antibody (Fig 9C), however the difference was not statistically significant. Overall inflammation was similar in the lungs of RV/PR8-infected mice treated with anti-IFNAR and control antibodies (Fig 9D). However, immunohistochemical (IHC) staining for a neutrophil-specific protein, Ly6G, indicated that IFNAR signaling limits neutrophilic inflammation in RV/PR8-infected lungs (Fig 9E). Enhanced neutrophilic inflammation in the lung parenchyma of anti-IFNAR treated mice was accompanied by increased infiltration of red blood cells into the alveoli (pulmonary hemorrhage) on day 6 (Fig 9D). Altogether, this suggests that type I IFN signaling plays a role in reducing, but not eliminating, viral replication and curtailing both neutrophil spread within the lungs and pulmonary hemorrhage in RV/PR8-infected mice.

In order to evaluate RV-induced recruitment of cells in the airways of mice, we quantified macrophages, neutrophils and lymphocytes in BALF from mice that received mock or RV with or without anti-IFNAR treatment on day 2 after inoculation (day 0 of study). Inoculation with RV did not increase the overall cell counts in BALF, but dramatically changed the composition from predominantly macrophages to neutrophils (Fig 10). Inhibition of IFNAR signaling did not alter cellular recruitment in response to RV infection.

**Fig 10.**
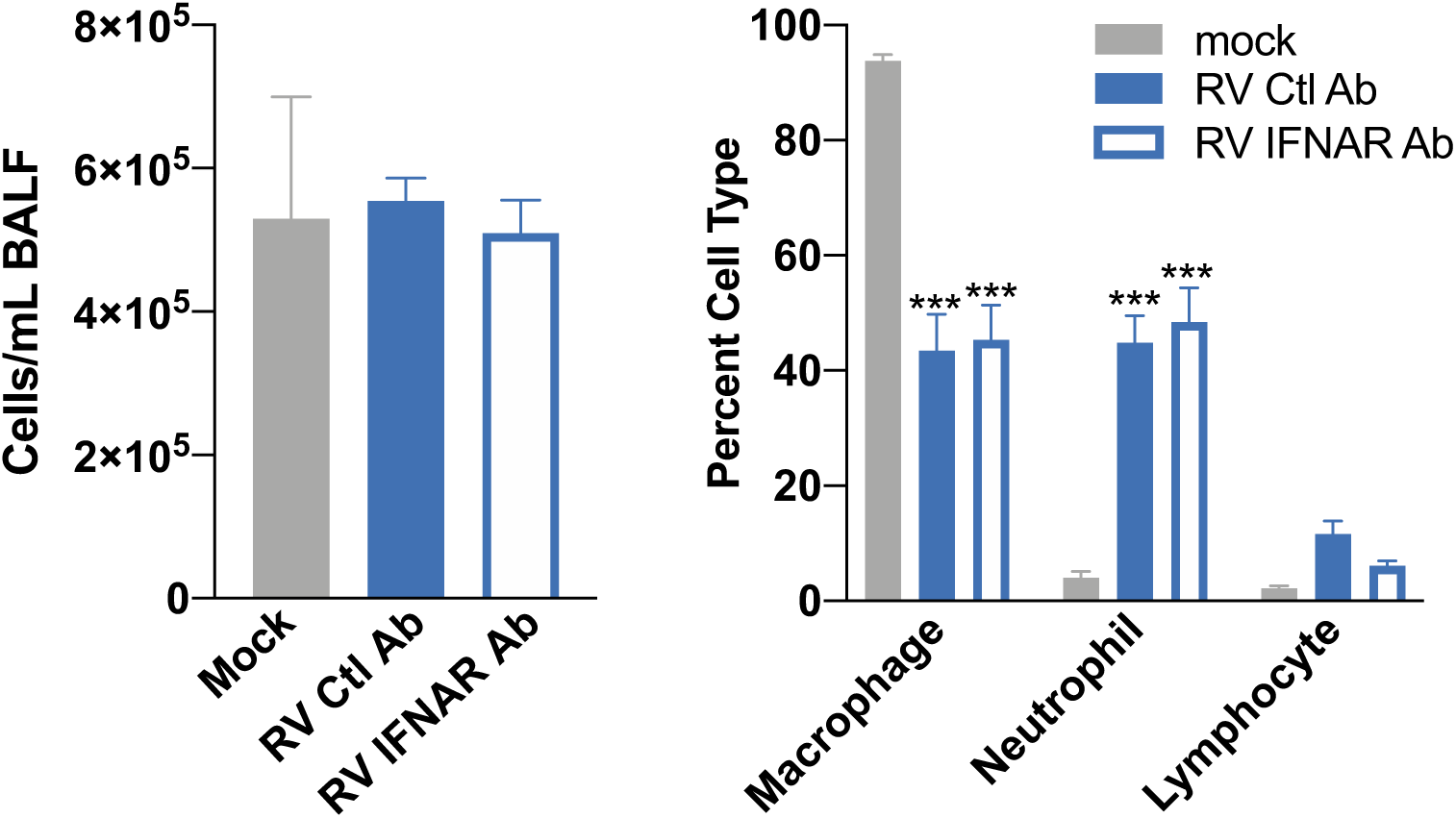
Airway cellular response to RV inoculation. Cytospins from BALF were stained with HEMA 3 to identify airway macrophages, neutrophils, and lymphocytes. At least 300 total cells were counted from duplicate cytospins for each sample to determine the percentage of each cell type. Total cells were counted in the BALF samples prior to cytospin preparation. Data shown are means and standard errors from 5 mice per group. Statistical significance between groups was determined using unpaired t-tests corrected for multiple comparisons using the Holm-Sidak method, ***p<0.001.

## Discussion

Previously, we found that inoculation of mice with a mild respiratory viral pathogen, RV or murine coronavirus MHV-1, two days before PR8 provided significant protection against PR8-mediated disease (10). In this study, we expanded these results to show that RV-mediated protection was not specific to PR8, but also provided significant disease protection against a respiratory virus from another viral family, PVM. This is in agreement with other studies showing heterologous immunity by respiratory viral pathogens in mouse models (11–13);(14). Recently, RV has been shown to inhibit replication of influenza A virus and SARS-coronavirus-2 in cultured airway epithelial cells (9, 49). Interference by respiratory viruses within a human host may contribute to reduced disease severity and alter population-level viral dynamics. Coinfection by rhinoviruses has been associated with reduced severity of the 2009 pandemic influenza A virus strain despite similar H1N1 viral loads (50). Conversely, infection with influenza viruses has also been found to reduce the severity of rhinovirus infections (51). Epidemiological studies have suggested that co-circulating viruses, especially rhinoviruses, can inhibit population-level dynamics of other respiratory viruses (49, 50, 52–54). In 2009, introduction of the new pandemic H1N1 strain delayed the circulation of other seasonal respiratory viruses (55). A large study of children with community acquired pneumonia found the incidence of rhinoviruses to be high, however rhinoviruses were found less frequently in combination with other RNA viruses, including influenza and RSV, than would be expected if coinfection were random (3). Similar findings have been reported in adult populations (49). Another large, multi-center study of infants observed significant interference between RSV and RV (56). A prospective study of children found that infection with RV did not affect the likelihood of having an influenza infection the following week, but an influenza infection decreased the chances of having RV the following week (57). In a study of RSV and influenza cases across seven seasons in Israel, they found that when the peak of RSV cases coincided with the influenza peak, the percent RSV positive cases were lower than when the RSV cases peaked prior to influenza (14). While these studies suggest an important role of respiratory viral coinfection in viral pathogenesis and epidemiology, animal models are necessary to understand the interactions between co-infecting viruses and their host that may contribute to these observations.

Despite the commonality of RV reducing disease severity of two heterologous lethal viruses in our mouse model, there are differences between the virus combinations in the kinetics of disease and viral replication. Inoculation of mice with RV provided more effective protection against PVM than PR8. RV/PVM infected mice had little to no signs of disease and significantly limited PVM gene expression. In contrast, coinfection by RV prevented mortality, but not morbidity, associated with PR8 infection, and reduced viral gene expression but did not prevent infection by PR8 (10). Further, RV given concurrently with PVM was as effective as when it was given two days before PVM. In contrast, we previously showed that RV was less effective at reducing the severity of PR8 when given concurrently and also exacerbated disease when it was given two days after PR8 (10). We also observed differences in the kinetics of gene expression in response to these virus pairs. Host and viral gene expression changes in response to PVM were delayed compared to PR8, thereby giving a larger window for RV-mediated protection. Thus, RV may be inducing antiviral mechanisms that are more effective against PVM, or different mechanisms may be responsible for inhibiting PVM infection and mediating effective clearance of PR8. In support of the latter, we found that IFNAR signaling was required for RV-mediated reduction in disease severity during PR8, but not PVM, infection.

While we do not know if RV replicated in the airways or lungs, intranasal inoculation with a high dose of RV resulted in dramatic up-regulation of host gene expression prior to inoculation with the second, lethal virus. Despite expression of several chemokine and chemokine receptor genes, we did not observe a dramatic increase in immune cells in the lungs of RV-inoculated mice on day 0. Our flow cytometry panel was limited to focus on inflammatory cells (neutrophils and macrophages) and T cells and likely missed other cell types that could be recruited by RV early in infection. Further, by using whole lung for our assays, we may miss populations of cells that are small, but significant, in the airways. Indeed, other studies have found that neutrophils and lymphocytes are increased in the airways of RV-infected mice two days after inoculation (39, 58), which we also detected by analyzing BALF. While the chemokine signaling genes we identified in RV-inoculated mice on day 0 were also increased in single virus infected mice later, early upregulation of immune cell recruitment in RV-treated mice may contribute to early control of infection and reduced disease severity.

In contrast to early recruitment of neutrophils, RV-treated mice had reduced numbers of neutrophils and interstitial macrophages later during infection, which could be a result of reduced viral infection and/or direct down-regulation of the inflammatory response. Other studies have shown that rhinoviruses inhibit macrophage responses to bacterial infection and toll-like receptor (TLR) stimulation (59, 60). This down-regulation leads to reduced neutrophil recruitment and activation and results in enhanced bacterial infection. While suppression of TLR signaling is detrimental during a subsequent bacterial infection, it can be protective in the case of respiratory viral infections for which inflammatory responses contribute to disease pathology. Blocking signaling from multiple TLRs (TLR2, TLR4, TLR7, and TLR9) through inhibiting the adaptor protein TIRAP reduces the severity of a lethal PR8 infection in mice (61). Inhibition of TIRAP reduces PR8-induced cytokine production by macrophages and is likely protective by limiting inflammatory responses (61). Similarly, inhibition of TLR2 and TLR4 signaling during PR8 infection reduces inflammatory cytokine responses and disease severity (62).

Inflammatory responses, particularly by granulocytes, are also associated with disease pathogenesis of PVM infection, independent of viral replication (34, 35). Priming with probiotic bacteria or parasitic infections can reduce the severity of PVM, which corresponds with reduced pulmonary inflammation. When given intranasally on days -14 and -7 prior to, or days 1 and 2 after PVM inoculation, *Lactobacillus spp.* prevent lethal viral infection (28, 63, 64). Bacterial priming is associated with reduced production of proinflammatory cytokines and chemokines and inflammatory cell recruitment to the lungs, especially neutrophils (28, 63, 64). Similarly, chronic schistosomiasis protects mice from a lethal PVM infection, which corresponds with up-regulation of mucus secretion pathways (32).

We found that RV-treated mice had increased expression of several genes in the mucin production pathway. Muc5ac, the predominant gel-forming mucin in the airways of humans and mice, is produced by goblet cells in the airway epithelium and Clca1 is a chloride channel regulator that is needed for proper mucus hydration. The forkhead transcription factor, FoxA3, induces expression of multiple genes in the mucus production pathway that we found to be concurrently upregulated, including Muc5ac, Muc5b, Agr2, and Itln1 (65). Chen et al. demonstrated increased expression of FoxA3 and Muc5ac and goblet cell metaplasia in the airway epithelium of mice 3 days after infection by RV (65), and additional studies have reported up-regulation of Muc5ac mRNA and protein in the lungs and airways of RV-infected mice (39, 65, 66). Influenza A virus strains, including PR8, also induce expression of Muc5ac and mucus secretion (67–69). While stimulation of mucus production can promote pathology in chronic airway diseases such as asthma and COPD, acute mucus production in hosts with healthy airways promotes innate defense against pathogen invasion. Over-expression of Muc5ac in mice has been shown to reduce PR8 infection and neutrophil recruitment (33). Muc5ac also promotes neutrophil transmigration and recruitment (70), thus the reduction in neutrophils seen by Ehre *et al*. may be a consequence of lower viral replication. Muc5ac has also been shown to reduce the severity of PR8 and PVM infections in a parasite coinfection model (32). Production of Muc5ac is induced by multiple inflammatory mediators, including IL-4, IL-13, EGF, and TGF-*α*, and is enhanced by additional signaling molecules through MAPK-dependent pathways (71). Muc5ac is also upregulated by IFN-β in human bronchial and alveolar epithelial cells and mouse lung tissues, and is dependent on signaling through aryl hydrocarbon receptor (AhR) (72). Thus, upregulation of Muc5ac by RV is likely not prevented by inhibition of IFNAR signaling and thus not sufficient to prevent PR8 lethality. Future studies will address the potential role of mucins in RV-mediated protection against PVM disease.

We found dramatic differences in the kinetics of interferon (IFN)-response gene expression in PR8-vs. PVM-infected mice, which corresponded with differences in the requirement of IFNAR signaling in protection against PR8, but not PVM. Although PVM encodes two nonstructural proteins with potent antagonist activity against type I IFN, treatment of mice with type I IFN prior to PVM infection is protective (22). It is possible that the IFN response to RV is not robust enough to account for protection against PVM or multiple, redundant mechanisms may contribute to protection, for example type I and type III IFN (22).

Type I IFN signaling is important for immediate antiviral defense mechanisms as well as orchestrating the correct balance of immune cells responding to infection. Others have shown that type I IFN signaling is critical for orchestrating monocyte responses to PR8 infection, which in turn limits neutrophilic inflammation (20). We observed increased numbers of neutrophils in BALF and through-out lung tissues in RV/PR8 coinfected mice that were treated with an IFNAR inhibiting antibody. These findings demonstrate that type I IFN can provide protection during respiratory viral infections that is independent of direct inhibition of viral replication and correlates with reduced neutrophilic inflammation and tissue damage. Type I IFN responses are also important in the activation of T cell-mediated immune responses (73–75). While they did not reach statistical significance, the numbers of CD4+ and CD8+ T cells in the lungs were modestly increased in RV/PR8-compared to mock/PR8-infected mice on day 6. Future studies to evaluate potential functional differences in lung-resident and recruited T cell subsets and their roles in protection will be critical for a complete understanding the type I IFN-dependent mechanisms whereby RV reduces the severity of PR8 infection.

## Materials and Methods

### Ethics Statement

All mouse procedures were approved by the University of Idaho’s Institutional Animal Care and Use Committee, in compliance with the NIH Guide for the Care and Use of Laboratory Animals. Female, 6-7 week-old BALB/c mice were ordered from Invivogen/Envigo and were allowed to acclimatize for 10 days prior to experimentation. All mice were housed in the UI Laboratory Animal Research Facility under 12 hour light/dark cycles, received food and water *ad libitum*, and were monitored daily for any signs of distress. Mice were humanely euthanized if they reached endpoints including more than 25% weight loss and/or severe clinical signs of disease.

### Virus Infections

Viruses used in this study include PVM strain 15 (ATCC VR-25), RV1B strain B632 (ATCC VR-1645) and influenza A virus, A/Puerto Rico/8/34 (H1N1; BEI Resources NR-3169). Viruses were grown and titrated by TCID_50_ assays in BHK-21 (PVM), HeLa (RV1B), and MDCK (PR8) cell lines.

Mice were anesthetized using isoflurane during intranasal inoculation. RV/PVM timing experiments were performed using groups of seven mice. Mice were either mock-inoculated (PBS/2% FBS) or inoculated with 7.6 x 10^6^ TCID_50_ units of RV1B two days before, simultaneously with, or two days after 1.0 x 10^4^ TCID_50_ units PVM in 0.05 mL intranasally. Mice were then monitored daily for mortality, weight loss, and clinical signs of disease (ruffled fur, hunched posture, lethargy, and labored breathing). Clinical signs in these four categories were scored on a scale of 0-3 (0-none, 3-severe). Humane euthanasia was performed when mice lost more than 25% of their starting weight or exhibited severe clinical signs of disease. Survival and weight loss data were analyzed with Prism 8.0 (Graphpad) software using Log-rank Mantel-Cox and student’s *t*-test corrected for multiple comparisons using the Holm-Sidak method, respectively.

RNA-seq and flow cytometry experiments were performed using groups of five mice. Mice were either mock-inoculated (PBS/2% FBS) or inoculated with 7.6 x 10^6^ TCID_50_ units RV1B intranasally on day -2. On day 0, mice were either inoculated with ∼50 TCID_50_ units of PR8 or 1.0 x 10^4^ TCID_50_ units of PVM. Mice were euthanized on days 0, 2, 4, and 6 to collect lungs for analyses. Left lobes were used for flow cytometry analysis and right lobes were placed in RNALater for RNA-seq analysis (see below). We consistently see infection in right and left lobes upon intranasal inoculation of PR8 or PVM in 50 uL volumes (S11 Fig).

Signaling by interferon alpha receptor 1 (IFNAR1) was inhibited by treating mice with 0.05 mg anti-IFNAR1 antibody (MAR1-5A3, Bio X Cell) intranasally on days -2 and 0 with virus or mock inoculations. Antibody of the same isotype (mouse IgG1k, MOPC-21, Bio X Cell) was used as a negative control. Mice were monitored and weighed daily as described above and groups of 5 mice were euthanized at pre-determined time points to quantify viral replication, IFN-induced gene expression, and cellular infiltration.

### RNA-seq Analysis

RNA was extracted from mouse lung tissue according to the RNeasy Plus with gDNA removal protocol (QIAGEN) and quantified using an HS RNA kit and Fragment Analyzer (Advanced Analytical). The three samples with the highest RNA quality from each group were used for RNA-seq analysis. Stranded RNA libraries were prepared from 4 ug RNA of each sample by UI’s IBEST Genomics Resources Core according to KAPA’s stranded mRNA-seq (KK8420) library preparation protocol with capture of polyadenylated mRNAs. Libraries were tagged with unique ligation adapters (BioOScientific), amplified, quantified (Qubit and AATI Fragment Analyzer), and pooled at an equimolar ratio. The pooled library was split, pooled with other libraries, and sequenced across 5 lanes of an Illumina HiSeq4000 100bp PE run at University of Oregon’s Genomics and Cell Characterization Core. Paired end reads (100 bp) were quality trimmed and filtered using Trimmomatic v0.36 (ILLUMINACLIP:2:20:10, HEADCROP:10, SLIDINGWINDOW:4:15, MINLEN:36) and mapped to the mouse genome (GRCm38/mm10 downloaded from UCSC) using TopHat v2.1.1 (r 300) (76, 77). The TopHat alignments were the starting points for two methods of quantifying read counts and four methods of analyzing differential expression. The Cufflinks v2.1.1 package (Cufflinks, Cuffmerge, Cuffdiff) was compared to DE pipelines using HTSeq v0.8.0 (htseq-count -m intersection -nonempty -s reverse -t exon) followed by DESeq2 v1.18.1, and EdgeR 3.18.1 (78, 79). DESeq2 and EdgeR were run within the SARTools wrapper (80). DEGs with adjusted p values below 0.05 were dropped. A text file containing important parameters and code is included in S1 File.

#### Analysis of viral gene expression

Reads that did not map to the mouse genome were extracted and de-interleaved using BBmaps’ reformat script (http://sourceforge.net/projects/bbmap/) to give pair reads for mapping. These reads were aligned to the RV1B (D00239), PR8 (LC120388-LC120395), and PVM (AY729016) genomes using TopHat v2.1.1 and gene coverages were counted using HTseq v0.8.0 (see parameters above). Statistical significance was determined using a generalized linear model in R.

#### DEG function analysis

The DEG lists for each treatment comparison were searched for differentially enriched gene ontologies (GO terms) and KEGG pathways using CompGO (81). CompGO utilizes gene expression level data (as opposed to just gene names) to identify potential functions that differ between treatments. We ran CompGO on DEG lists from each individual method (Cufflinks, EdgeR, DESeq2) and combinations of gene sets to anecdotally look for consistency across methods.

### RT-qPCR

RNA was extracted from lung tissues using Trizol (Invitrogen) or RNAeasy Plus (QIAGEN) reagents following the manufacturers’ protocols. cDNA synthesis was performed using SuperScript IV VILO master mix (Invitrogen) and qPCR was done with PowerUp SYBR Green (Applied Biosystems) using a StepOnePlus real-time instrument (applied Biosystems). Published primer sets were used to quantify PR8 HA (82), PVM SH (83), IFN-β (84), Mx1 (73), GAPDH (85), and β-actin (86). Mx1 and IFN-β C_T_ values were normalized to GAPDH and the resulting ΔC_T_ values were normalized to those from mock-inoculated mice and expressed as 2^-ΔΔCT^. C_T_ values of viral genes were normalized to β-actin and expressed as 2^-ΔCT^.

### Flow Cytometry

The left lung lobe of each sample was dissociated in RPMI medium containing type IV collagenase (MP Biomedicals) and DNase I (Spectrum) using gentleMACS C tubes according to the manufacturer’s recommendations (Miltenyi Biotec). Cells were filtered through a 70 µm strainer and red blood cells were lysed before blocking F_C_ receptors using anti-CD16/CD32 (BioLegend). Cell-specific proteins were labeled with the following antibodies: CD11b-Alexa Fluor 488 (eBiosciences), Ly6G-APC (eBiosciences), CD64-PE (BioLegend), SiglecF-PerCP-Cy5.5 (BD Biosciences), or the appropriate isotype control antibodies. T cell subsets were stained with antibodies against CD3-PerCP-Cy5.5 (eBiosciences), CD8-APC (eBiosciences), and CD4-Alexa Fluor 488 (BioLegend). Stained cells were incubated in BD Stabilizing Fixative (BD Biosciences) and analyzed using a FACS Aria Cytometer (BD Biosciences). Results were analyzed using FlowJo software (Treestar) and gating was performed based on fluorescence-minus-one controls. The gating strategy used to identify neutrophils, alveolar macrophages, and interstitial macrophages is shown in S8 Fig. T cells were gated as CD3+/CD4+ T-helper cells and CD3+/CD8+ cytotoxic T cells.

### Quantification of Viral Loads

Infectious virus was quantified in homogenized lungs or BALF samples by TCID_50_ assays in HeLa (RV), MDCK (PR8) and BHK21 (PVM) cell lines. We have previously found that the presence of RV in samples does not interfere with titration of PR8 or PVM in MDCK cells or BHK21 cells, respectively (10).

### Analysis of BALF cells

BALF was collected by tracheal cannulation and lavage with 1 mL PBS twice per sample. Collected cells were counted and spun onto microscope slides in duplicate samples, which were stained with Hema 3 staining solutions (Fisher Scientific). Gridded coverslips and microscopy were used to count at least 300 neutrophils, macrophages and lymphocytes per sample and these counts were used to calculate the percentage of each cell type.

### Histology and Immunohistochemistry

Lung tissues from two animals per treatment group and time point were processed and stained as previously described (10). Antibodies specific for PR8 hemagglutinin protein (NR-3148, BEI Resources) and Ly6G (1A8, Bio X Cell) were detected with immPACT vector red and immPACT DAB, respectively (Vector Laboratories).

### Statistical Analyses

We analyzed flow cytometry (Fig 7) and viral read count (Fig 4) data resulting from our experiments to identify time-varying differences between mice infected with PR8 or PVM alone or pre-treated with RV. To this end, we used negative binomial regression on each response variable with an explanatory model that had a main effect of days post-infections, a main effect of treatment, and the interaction between the two main effects. Response variables were the number of CD11b cells, neutrophils, interstitial macrophages, alveolar macrophages, and viral read counts; all response variables were normalized versus total cell count except for viral RNA read count which was normalized against total RNA read count. Due to *a priori* visual investigation of our data, it seemed some of our response variables might be better fit with a quadratic time term. Because of this we fit an alternative model that included an orthogonal polynomial of degree 2 for time. We assessed whether the quadratic model was better than the linear model using a likelihood ratio test and chose the quadratic model if it offered a significant improvement in fit over the simpler linear model. The significance of treatment and time was determined using a type-I ANOVA. To detect differences between treatments at a given time, we also performed *post hoc* pairwise comparisons of the modeled mean at our observational time points (days 0, 2, 4, and 6) using the emmeans package in R. Supplemental Table 3 provides information on the analyses, including which time model was used, the significance of treatment and time. Significant contrasts are shown in individual figures.

Analysis of survival curves, weight loss, viral titers, RT-qPCR, and BALF cell count data were analyzed with Prism 8.0 (Graphpad) software using the methods described in each figure legend.

## Data Availability

Raw Illumina reads have been deposited in NCBI SRA database under accession numbers SRR7060120-SRR7060117 under project PRJNA453386.

## Acknowledgements

This study was supported by the National Institutes of Health (NIH) grants P20 GM104420 and P20 GM103397. Maricris Naranjo was supported by the REU Summer Program in Molecular and Organismal Evolution, funded by the National Science Foundation (Award #1757826). We thank the Institute for Bioinformatic and Evolutionary Studies (IBEST) Genomics Resources Core and Optical Imaging Core and the laboratory of Dr. Shirley Luckhart for support. We would also like to acknowledge the excellent technical support of Dr. Bhim Thapa and Mr. Kevin Hutchison. The following reagents were obtained through the NIH Biodefense and Emerging Infections Research Resources Repository, NIAID, NIH: Influenza A Virus, A/Puerto Rico/8/34 (H1N1).

## Supporting Information

**Supplemental Table 1** provides raw read counts (mean of the three replicates and normalized to library size) and log2 fold change values for genes that had statistically significant differences in expression between each comparison.

**Supplemental Table 2** provides statistically significant GO terms for each comparison.

**Supplemental Table 3** provides a summary of results from negative binomial regression analyses of viral read count and flow cytometry data.

**S1 Fig. Weight loss of mice used in the RNA-seq and flow cytometry experiments.** Mice were inoculated intranasally with saline (mock) or RV on day -2 and PR8 or PVM on day 0. Weight was recorded daily for all mice until the pre-determined time points. Days -2 through 2, N=15 per group; days 3 through 4, N=10 per group; days 5 through 6, N= 5 per group.

**S2 Fig. Read coverage of individual viral genes.** The proportion of reads mapping to each viral gene is shown for mice infected with PR8 or PVM.

**S3 Fig. Heatmap of mock/PR8 vs. RV/PR8 unique genes.** Expression of genes that are significantly different between mock/PR8 and RV/PR8 infected mice that are not significant between mock/PVM and RV/PVM infected mice. Colors represent Z-scores for individual genes across infections and time points as described for Fig 6.

**S4 Fig. Heatmap of mock/PVM vs. RV/PVM unique genes.** Expression of genes that are significantly different between mock/PVM and RV/PVM infected mice that are not significant between mock/PR8 and RV/PR8 infected mice. Colors represent Z-scores for individual genes across infections and time points as described for Fig 6.

**S5 Fig. Heatmap of differentially expressed genes shared by RV-treated mice.** Expression of genes that are significantly different between mock/PVM and RV/PVM infected mice that are also significant between mock/PR8 and RV/PR8 infected mice. Colors represent Z-scores for individual genes across infections and time points as described for Fig 6.

**S6 Fig. Heatmaps of MSigDB Hallmark Interferon Alpha Response gene set shown in** Figure 6**, including gene names.** See Fig 6 legend for details.

**S7 Fig. Heatmaps of MSigDB Hallmark Inflammatory Response gene set shown in** Figure 6**, including gene names.** See Fig 6 legend for details.

**S8 Fig. Total cell counts and gating strategy for flow cytometry.** Left lungs were processed for flow cytometry staining with antibodies against CD11b, Ly6G, CD64, and Siglec F. Total cell counts for all samples and representative plots showing the gating strategy to identify neutrophils, alveolar macrophages, and interstitial macrophages are shown.

**S9 Fig. RNA-seq data showing chemokine gene expression related to innate inflammatory cell recruitment.** Expression of neutrophil and monocyte/macrophage specific chemokine genes from the RNA-seq analysis are shown as Log2 fold change vs. mock-inoculated mice.

**S10 Fig. Anti-IFNAR treatment of RV-inoculated mice.** Nine mice per group were inoculated intranasally with 7.6 x 10^6^ TCID_50_ units of RV on day -2 and received a mock inoculation of buffer on day 0. Mice received either anti-IFNAR or control antibody with the inoculations on days -2 and 0. Five mice per group were monitored daily for survival and weight loss. RV was titrated from homogenized lung and BALF samples on days -1 and 0 from 4 mice per group by TCID_50_ assay.

**S11 Fig. Comparison of infection levels in right and left lobes.** Mice were inoculated with mock/PR8 or RV/PR8 as described in materials and methods. RT-qPCR was performed on RNA isolated from right or left lobes of individual mice on day 4 after PR8 inoculation. Microscopic images are shown from serial imaging of lung tissue sections from a mock/PR8-infected mice on day 4 of infection stained for HA antigen visualized with impact red. Images are representative of at least 3 mice.

